# Structure-Guided Computational Analysis of Linker effects in an scFv Targeting Guanylyl Cyclase C

**DOI:** 10.64898/2026.03.30.714862

**Authors:** Teresa S. Viegas, Rita Melo

**Author notes:** Corresponding author (Rita Melo).

## Abstract

Single-chain variable fragments (scFvs) are widely used in diagnostic and therapeutic applications. These antibody fragments comprise two antibody variable domains connected by a flexible peptide linker whose properties critically influence folding, stability, oligomeric state, and antigen-binding. Therefore, careful linker selection represents a key step in scFv design. Guanylyl Cyclase C (GUCY2C) is a tumor-associated cell surface receptor expressed in gastrointestinal malignancies, including more than 90% of colorectal cancer (CRC) cases across all disease stages. Its restricted physiological expression pattern makes GUCY2C an attractive target for immunotherapy and precision oncology therapies. Here, we investigated the structural and functional consequences of incorporating alternative linker designs into an anti-GUCY2C scFv. Using molecular modeling, protein-protein docking, and molecular dynamics (MD) simulations, we evaluated the conformational stability, interdomain organization, and antigen-binding interactions of each construct. Our results provide a dynamic, structure-based assessment of how linker composition influences GUCY2C recognition and scFv structural behavior. Furthermore, this work establishes a computational framework for the rational optimization of GUCY2C-targeted antibody fragments.

## 1. Introduction

Single-chain variable fragments (scFvs) are widely used in diagnostic and therapeutic applications. The variable region of the antibody, comprising the variable light (V_L_) and variable heavy (V_H_) domains, constitutes the minimal structural unit required to preserve antigen recognition [1]. In scFvs, these domains are connected by a flexible peptide linker, whose length and sequence play a significant role in determining the expression levels, folding efficiency, oligomeric state, antigen affinity/specificity and solubility [2-4]. The glycine-serine (GS) repeat is the most common linker design owing to its flexible nature; however, it presents limitations, including challenges in PCR-based engineering and immunogenicity due to its repetitive nature [2, 5]. For these reasons, alternative non-repetitive designs have been proposed. These flexible linkers are also rich in GS but can incorporate polar amino acids such as lysine or glutamate to improve solubility, threonine or alanine to preserve flexibility, or proline to reduce proteolytic susceptibility [6-7]. The optimal linker for a given scFv construct is, however, dependent on the structural and functional context where it is inserted, thereby requiring thorough construct-specific evaluation [3, 8].

The intestinal epithelial receptor Guanylyl Cyclase C (GUCY2C) is normally expressed on the luminal surfaces of intestinal epithelium from the duodenum to rectum [9]. Its main function is the maintenance of intestinal homeostasis and is activated by the peptide hormones guanylin and uroguanylin, as well as the bacterial heat-stable enterotoxins produced by enterotoxigenic *E. coli* [10-11]. GUCY2C is widely expressed across CRC and other gastrointestinal tumors [12]. The loss of epithelial polarity and disruption of tight junctions in tumors may allow for the preferential uptake of GUCY2C-targeted therapies, making it a promising target for precision therapy approaches in CRC [13-14]. Recently, a T cell-engaging CD3 bispecific antibody targeting GUCY2C, PF-07062119, has shown strong preclinical activity. Specifically, PF-07062119 showed efficacy in multiple colorectal cancer models, including those harboring KRAS or BRAF mutations, which are associated with resistance to approved EGFR-targeted therapies [15].

Building on the therapeutic potential of GUCY2C, the scFv analyzed in this work corresponds to the GUCY2C-binding arm of PF-07062119. The crystal structure of this scFv in complex with the extracellular domain of GUCY2C has been previously resolved (Figure 1; PDB 8GHP), indicating that the scFv epitope is located on the receptor’s N-terminal H2 helix [16]. While this structure provides detailed insight into the recognition of GUCY2C by the antibody fragment, the dynamics of this interaction remain poorly characterized, and the potential influence of different linker sequences on binding behavior has not been systematically assessed.

**Figure 1:**
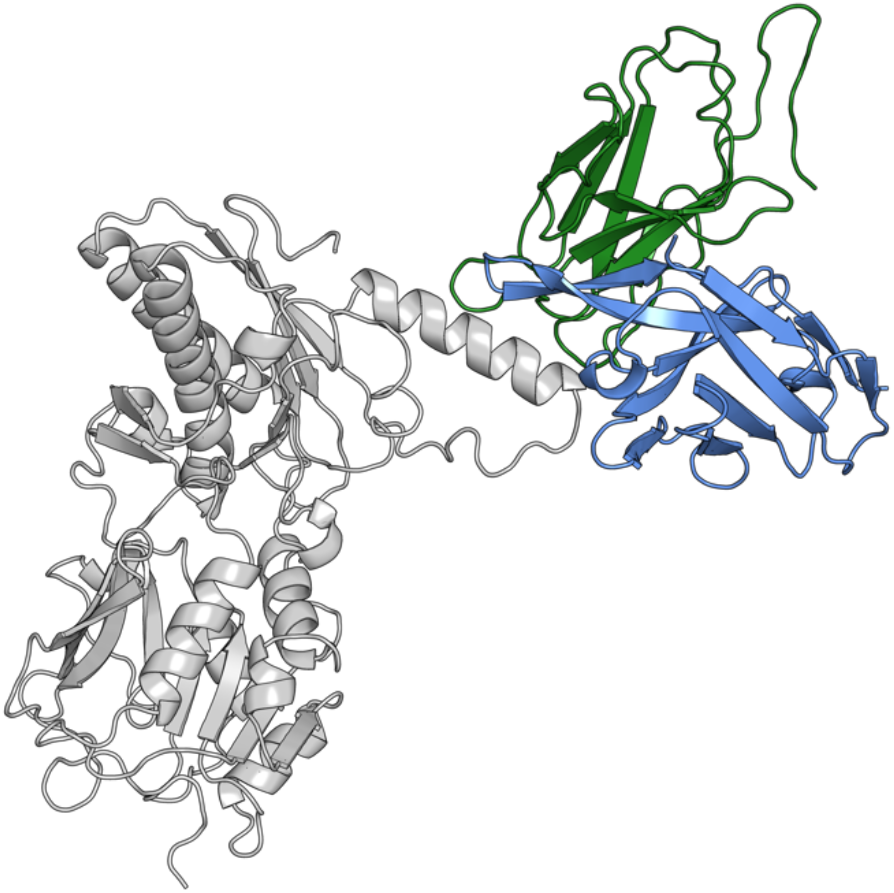
Depiction of the GUCY2C-ECD-anti-GUCY2C-scFv complex. GCC-ECD is colored in dark gray and the anti-GUCY2C scFv in blue (V_H_) and green (V_L_).

Here, we have employed a computational approach to investigate the dynamic behavior of the anti-GUCY2C-scFv-GUCY2C complex and to compare four alternative scFv designs incorporating different peptide linkers. Structural models of the scFvs in complex with GUCY2C were generated through protein structure prediction and docking, after which all-atom molecular dynamics (MD) simulations were performed to explore their overall conformational behavior. Binding free energy calculations and per-residue pairwise interaction analyses were subsequently performed to compare relative binding affinities and identify key interacting residues. This study provides a dynamic insight into the interaction of this anti-GUCY2C scFv with its target, and a computational framework to support the rational design and refinement of GUCY2C-targeted antibody constructs. Additional methodological details and supporting analyses are provided in the Supplementary Information.

## 2. Materials and Methods

### 2.1 Selection of Linkers

Linkers of varying lengths and compositions were selected from previously published studies on scFv linker design and optimization [6-7, 17]: L1 – GSTSGSGKPGSGEGSTKG; L2 – EGKSSGSGSESKST; L3 – GGGGSGGGGSGGGGS; L4 – GSAGSAAGSGEF.

### 2.2 Model Construction and Quality Assessment

Each scFv model comprises the V_L_ and V_H_ domains joined together by one of the selected linkers (L1-L4) in a V_L_-linker-V_H_ orientation. Amino acid sequences of both chains were obtained from Protein Data Bank (PDB) 8GHP. The 3D structures of the four anti-GUCY2C scFv models were predicted using the RoseTTAFold tool available in the Robetta webserver [18]. For each scFv, the five highest-confidence structures were retrieved and evaluated using MolProbity [19]. The MolProbity score was used to rank the five candidate structures of each scFv, from which the lowest-scoring ones were selected for molecular docking.

### 2.3 Molecular Docking

Protein-protein docking of each scFv with GUCY2C-ECD was carried out using the HADDOCK2.4 (High-Ambiguity-Driven protein-protein DOCKing) webserver [20, 21]. The GCC-ECD structure was retrieved from the anti-GCC-scFv-GCC-ECD antibody complex (Figure 1; PDB 8GHP). HADDOCK utilizes a three-stage docking and refining protocol, beginning with a full randomization of orientations and docking by rigid body energy minimization, followed by semi-flexible simulated annealing, during which the interface is considered flexible, and a final refinement through MD in explicit solvent [22, 23]. Biological information regarding the residues involved in the intermolecular interaction is incorporated through Ambiguous Interaction Restraints (AIRs), which drive the docking. We defined scFv active residues based on solvent accessibility analysis performed in PyMOL [24]. Residue-level changes in solvent-accessible surface area (SASA) were estimated by comparing each residue’s accessible surface area (ASA) in the complex to that of the isolated chains. Residues exhibiting a decrease in ASA greater than 1.0 Å^2^ were classified as interface residues and defined as active during docking. For GUCY2C, active residues included those within the experimentally defined PF-07062119 binding region identified by Rampuria et al. [16]. Among the 200 protein-protein complexes obtained in each run, the 10 top-scoring complexes, in terms of HADDOCK score, were retrieved for further analysis. From these, the final complex was selected based on the best balance between HADDOCK score and conservation of active residues at the interface.

### 2.4 Molecular Dynamics Simulations

Docked scFv-GUCY2C complexes and the free GUCY2C were subjected to all-atom molecular dynamics (MD) simulations in GROMACS version 2023.1 [25-27] using the AMBER ff99SB-ILDN force field [28]. The structures were solvated in a cubic TIP3P water box with 10 Å buffering distance to the box boundaries. Systems were neutralized and ionized with Na^+^/Cl^−^ ions to a final concentration of 0.15 M, and periodic boundary conditions were applied in all three dimensions. Energy minimization was performed using the steepest descent algorithm, followed by a two-step equilibration protocol: (i) 100 ps in the canonical (NVT) ensemble at 300 K using the velocity-rescaling (V-rescale) thermostat (τ_T_ coupling constant of 0.1 ps) [29]; and (ii) 100 ps in the isothermal-isobaric (NPT) ensemble at 1 bar using the Parrinello-Rahman barostat (τ_P_ = 2.0 ps, compressibility of 4.5 × 10–^5^ bar^−1^) [30-31]. Initial velocities were assigned according to a Maxwell-Boltzmann distribution at 300 K. Three independent production simulations per system were then carried out for 400 ns (scFv-GUCY2C complexes) and 300 ns (GUCY2C-only) in the NPT ensemble using a 2 fs integration timestep.

Temperature was maintained at 300 K using the V-rescale thermostat (τ_T_ = 0.1 ps), with separate coupling applied to protein and non-protein groups. Pressure was maintained at 1 bar under isotropic coupling conditions using the Parrinello-Rahman barostat (τ_P_ = 2 ps). Long-range electrostatic interactions were computed using the particle mesh Ewald (PME) method with a real-space cutoff of 1.0 nm, fourth-order spline interpolation, and a Fourier grid spacing of 0.16 nm [32-33]. van der Waals interactions were treated using a cut-off scheme with a 1.0 nm cutoff, and long-range dispersion corrections were applied to both energy and pressure. Neighbor lists were constructed using a Verlet cutoff scheme and updated every 20 steps. All bonds involving hydrogen atoms were constrained using the LINCS algorithm [34], enabling the use of a 2 fs timestep.

### 2.5 MD trajectories analysis

Protein structures were visualized using PyMOL. Trajectory analyses were performed using tools from the GROMACS suite, including calculation of root-mean-square deviation (RMSD), per-residue root-mean-square fluctuations (RMSF), and interdomain center-of-mass (COM) distances. The equilibrated portion of each trajectory was determined individually for each system and replicate based on the RMSD and interdomain COM distance profiles. RMSD calculations were performed using the backbone atoms of the full complexes (Figure S2). RMSF was calculated using only Cα atoms. COM distance plots were obtained by calculating the center-of-mass of GUCY2C-ECD and each scFv variable domain (Figure S3). Only frames corresponding to the equilibrated region were used for subsequent analyses.

Binding free energies of the scFv-GCC complexes were estimated using the MM/PBSA method, as implemented in gmx_MMPBSA [35]. Calculations were performed over 1500 evenly spaced frames extracted from the equilibrated region of each trajectory, following the single trajectory approximation. The AMBER ff99SB-ILDN force field was used to calculate the van der Waals (ΔE_vdW_) and electrostatic (ΔE_ele_) terms, and the level-set dielectric interface implementation (ipb = 2) was used in PB calculations. The energetic contribution of each individual interfacial residue (Table 1) was determined using the per-residue effective free energy decomposition protocol. The ionic strength was set to 0.15 M and all the other parameters were kept as default values.

**Table 1:**
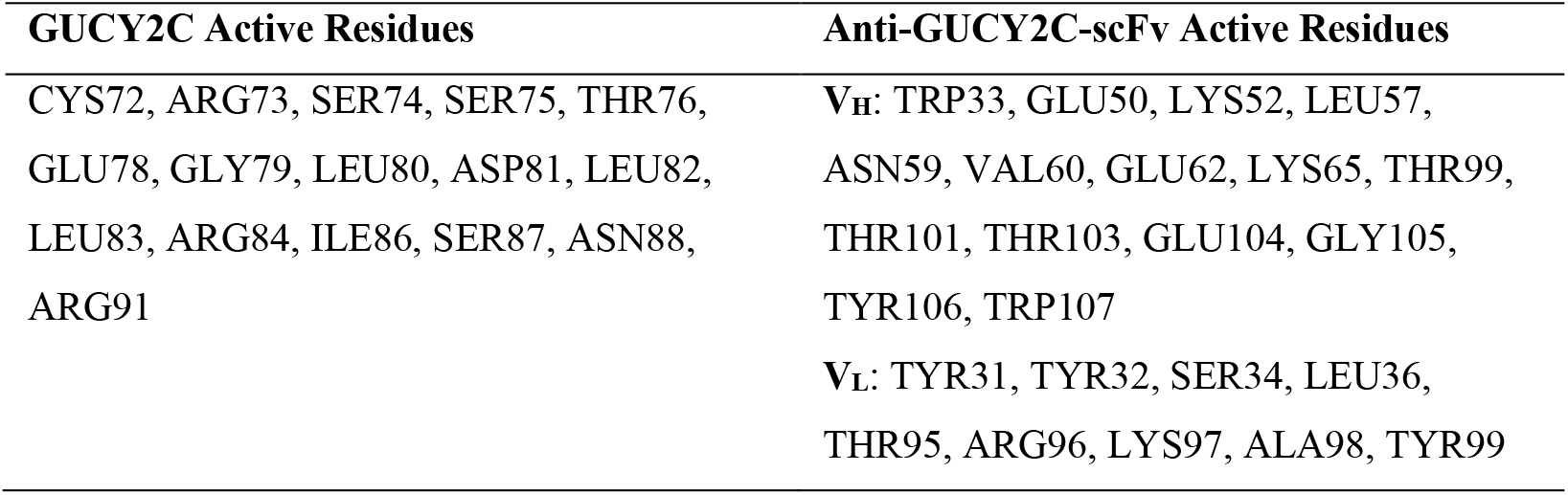
List of GUCY2C and anti-GUCY2C-scFv active residues used to guide docking.

| GUCY2C Active Residues | Anti-GUCY2C-scFv Active Residues |
| --- | --- |
| CYS72, ARG73, SER74, SER75, THR76,<br>GLU78, GLY79, LEU80, ASP81, LEU82,<br>LEU83, ARG84, ILE86, SER87, ASN88,<br>ARG91 | V<sub>H</sub>: TRP33, GLU50, LYS52, LEU57,<br>ASN59, VAL60, GLU62, LYS65, THR99,<br>THR101, THR103, GLU104, GLY105,<br>TYR106, TRP107<br>V<sub>L</sub>: TYR31, TYR32, SER34, LEU36,<br>THR95, ARG96, LYS97, ALA98, TYR99 |

Interprotein contacts were identified and quantified using an in-house script built upon MDTraj [36]. Additionally, PyInteraph2 [37] was used to identify interprotein hydrogen bonds, salt bridges, and hydrophobic contacts. Each interaction type was calculated using the default PyInteraph2 parameters.

## 3. Results

### 3.1 Model Construction and Docking

With no previously determined structure for the anti-GUCY2C-scFv bearing each linker, initial models were generated from the amino acid sequences using RoseTTAFold. For each linker, the five models with the highest predicted confidence scores were retrieved and superposed onto the experimentally determined scFv crystal structure to verify correct V_L_-V_H_ orientation and the overall conformation. Structural quality was subsequently assessed using MolProbity (Table S1, Supplementary Information), and the model with the best MolProbity score (lowest score) for each linker was selected as the final 3D structure (Figure S1, Supplementary Information).

To generate the models for each scFv bound to GUCY2C-ECD, protein-protein docking was carried out using HADDOCK. Docking was guided by both experimental and structural information. Previous epitope mapping studies identified residues 72-91 of GUCY2C as comprising the binding epitope of the anti-GUCY2C arm of PF-07062119; these residues were therefore defined as active residues on the receptor. On the scFv side, interfacial residues were identified from the GUCY2C-scFv crystal structure based on solvent-accessible surface area differences between the isolated chains and the complex and included as active residues to guide docking.

For each complex, the ten top-scoring docking solutions were analyzed. Final models were selected based on a combined assessment of HADDOCK score and conservation of active interface residues. This strategy prioritized docking conformations that both satisfied experimental restraints and exhibited favorable energetics. Docking statistics are summarized in Tables S3 and S4, and the selected representative complexes are shown in Figure 2.

**Figure 2:**
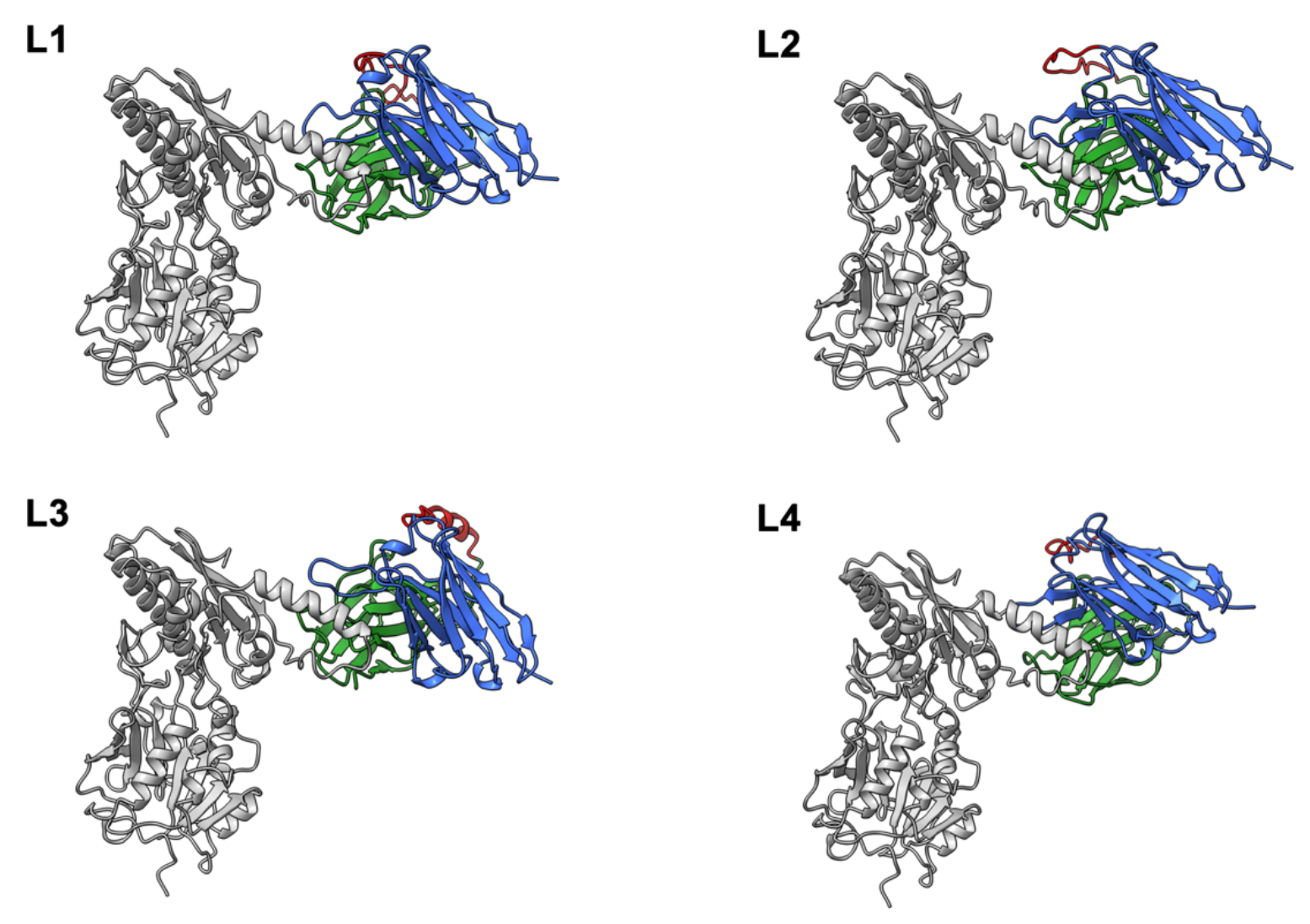
Best docking structure for each scFv model (L1-L4). GCC-ECD is colored in dark gray, the VL and VH domains of the scFv are colored in green and blue, respectively, and the linker in red.

### 3.2 Conformational Dynamics of the scFv-GUCY2C Complexes

To determine how the different linker sequences influence scFv-GUCY2C binding, and to obtain a detailed description of the interaction, atomistic MD simulations were performed for each scFv-GUCY2C complex. Equilibration was first assessed by monitoring both the backbone RMSD and the inter-domain COM distances for all scFv-GUCY2C systems and replicates (Figure S2 and S3). Equilibrium was considered reached once the running-average RMSD and COM distance profiles reached a stable plateau in each trajectory. For the GUCY2C-only simulations, equilibration was evaluated using RMSD alone (Figure S4). All subsequent analyses were performed over the equilibrated trajectory segments.

The RMSF of the Cα atoms was calculated to compare structural deviations on a per-residue basis across systems (Figure 3). On the scFv side, the linker region (residues 120-132/137, depending on linker length) exhibited the highest flexibility in all systems, consistent with its glycine- and serine-rich composition. Among the variants, linker L2 showed the greatest flexibility, whereas L1 displayed the lowest fluctuations across this region. While overall V_L_ and V_H_ domains exhibited similar fluctuation patterns across systems, some differences in magnitude were observed. In particular, the V_L_ domain of L2 displayed elevated fluctuations in loop regions spanning residues 30-33 and 70-73 compared with the other linker variants.

**Figure 3:**
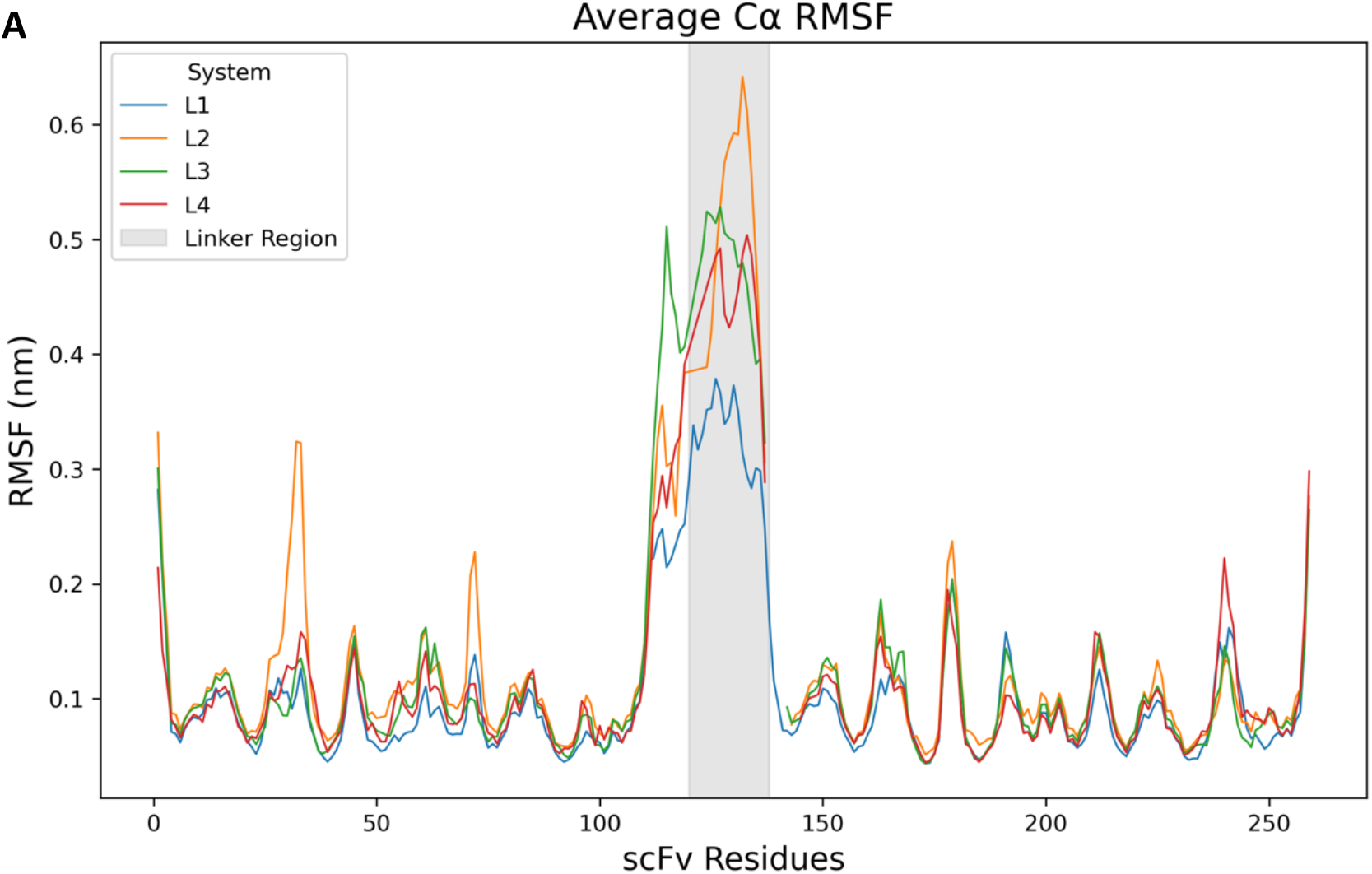
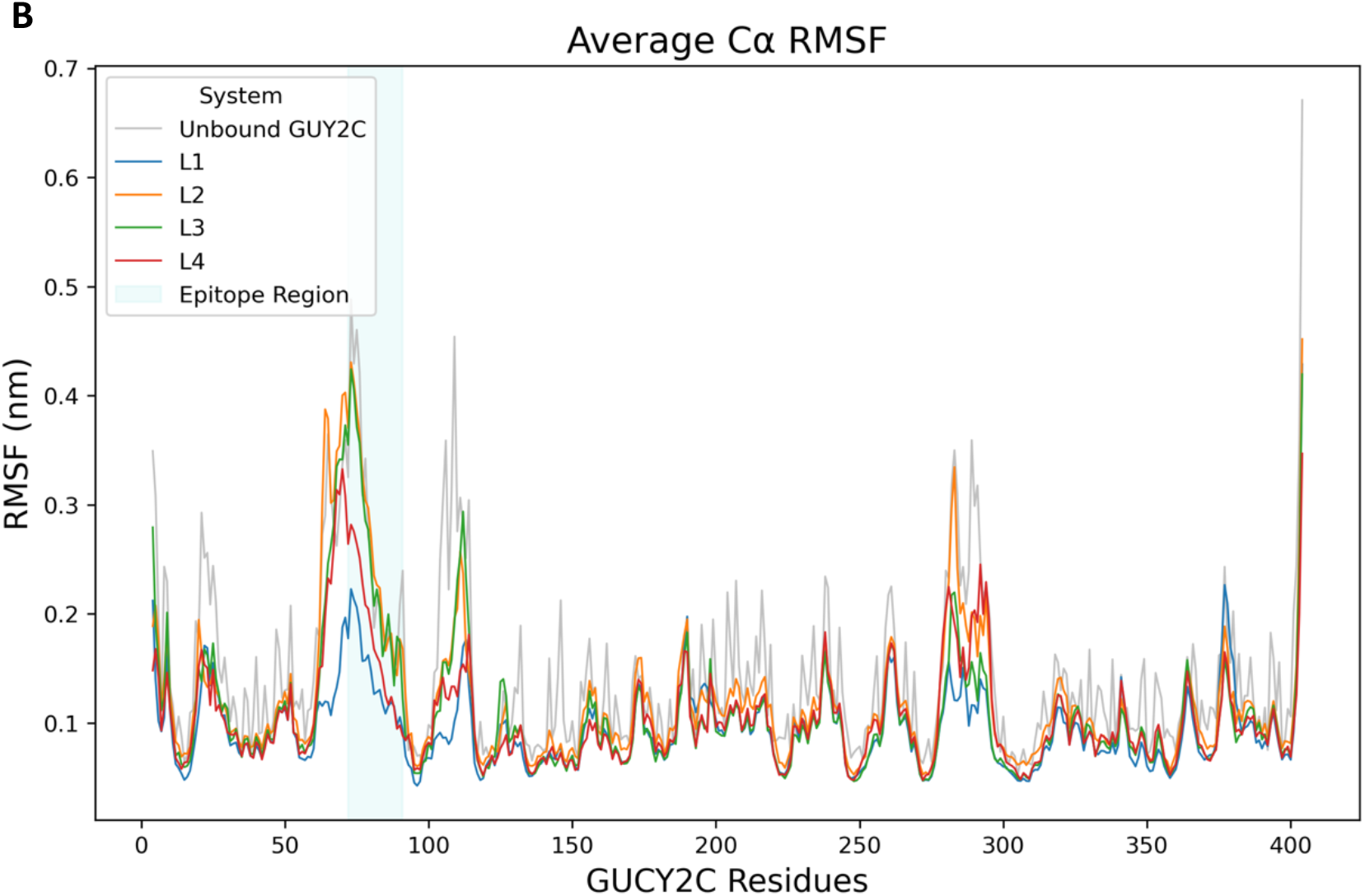
Average Cα RMSF values per residue for each scFv–GUCY2C complex (L1-L4), calculated over the equilibrated regions of the MD simulations. **A**) RMSF values per scFv residue; **B)** RMSF values per GUCY2C residue. Linker and epitope regions are indicated by the shaded area. RMSF values represent the average across three simulation replicates.

On the receptor side, the unbound GUCY2C exhibited higher overall RMSF values relative to the scFv-bound complexes, indicating that antibody binding generally reduces receptor flexibility. Within the epitope region (residues 72–91), distinct behaviors were observed across linker variants. L1 produced the lowest RMSF values within this region, indicating greater stabilization of the binding site. L4 induced moderate stabilization, whereas L2 and L3 displayed epitope fluctuations comparable to those of the unbound receptor. Beyond the epitope, two additional flexible regions in the unbound receptor (residues 100-115 and 275-300) showed reduced mobility in the L1 complex relative to the other systems, suggesting a broader stabilization of the receptor in the presence of the L1 scFv (Figure 4).

**Figure 4:**
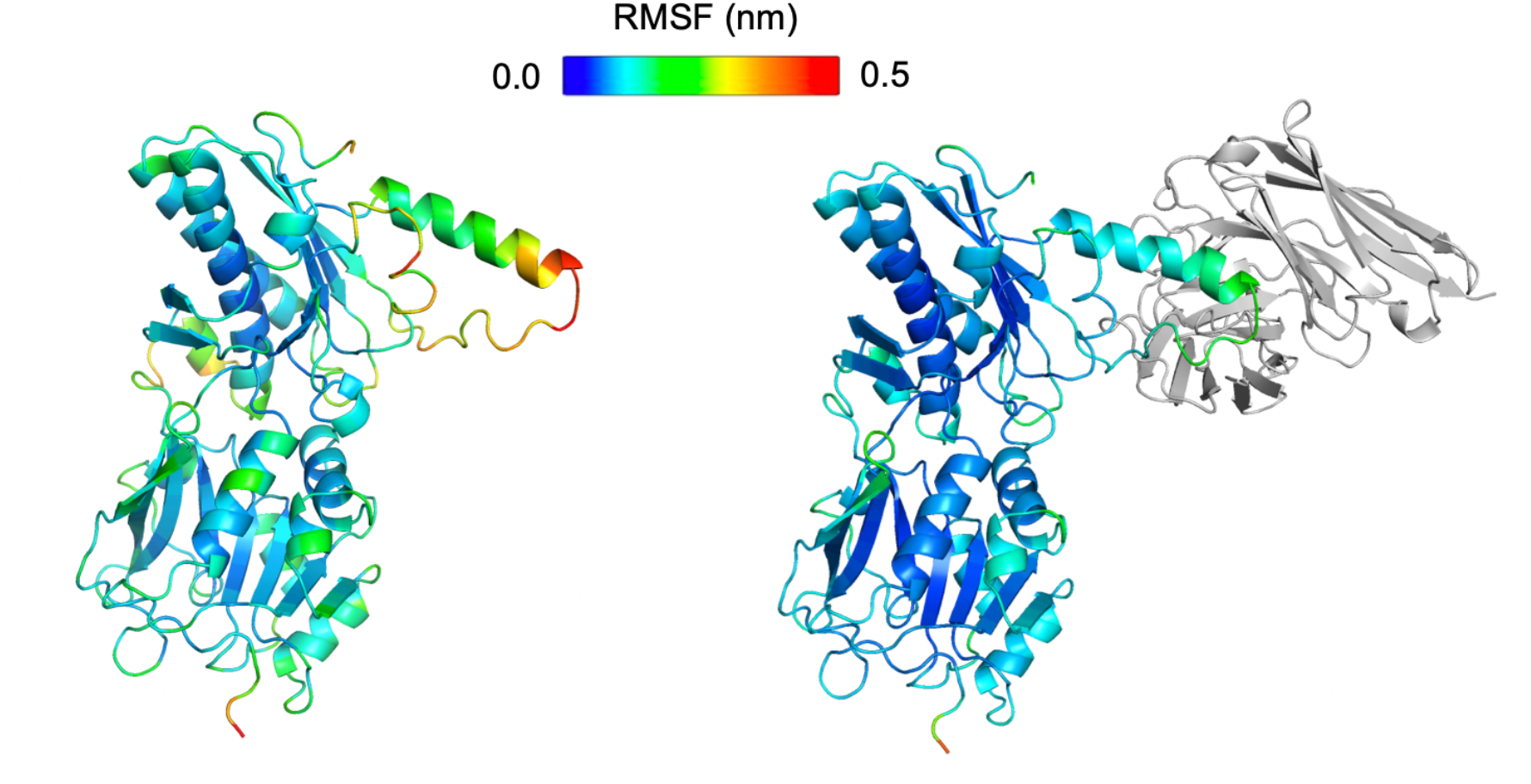
Structures of the unbound (left) and L1-scFv bound GUCY2C (right), colored by average Cα RMSF. Colors are distributed over the range of RMSF values, with dark blue representing the lowest, and red the highest RMSF.

**Figure 5:**
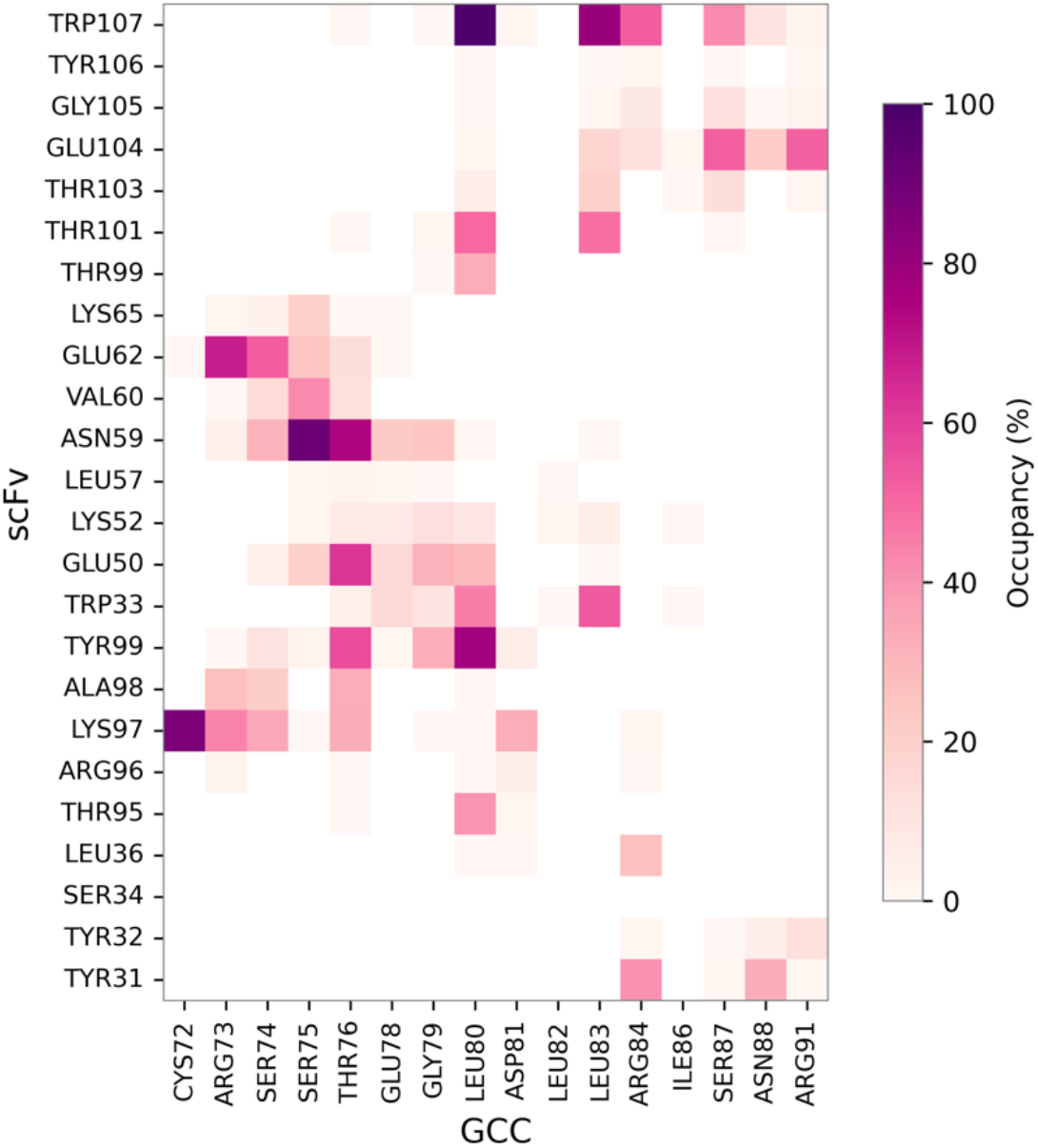
Two-dimensional inter-protein contact map of the L1 scFv-GUCY2C complex. The mean contact occupancy across three simulation replicates is shown, defined as the percentage of frames in which a given scFv-GUCY2C residue pair remained within 0.5 nm. Residues from GUCY2C are shown on the x-axis and scFv interface residues on the y-axis.

**Figure 6:**
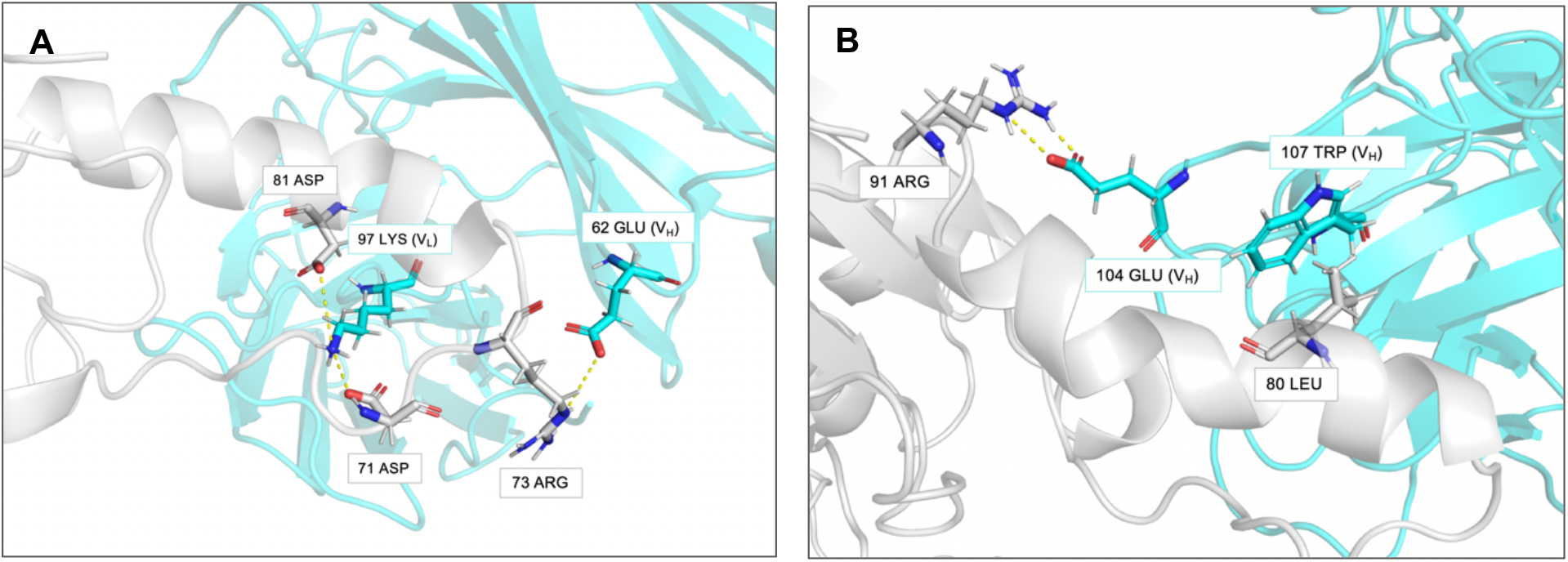
Representation of the interface between L1 scFv and GUCY2C. The scFv is shown in cyan cartoon and GUCY2C in white cartoon. Side chains of interacting residues are shown as sticks and colored by atom type and chain. Selected pairwise interactions are highlighted with yellow dashed lines. Panels A and B show representative structures from portions of the trajectory where the depicted interactions are most prevalent.

### 3.3 Binding Free Energy Profiles across Linkers

To compare the relative binding affinities of the scFv bearing different linkers, binding free energies of the scFv-GUCY2C complexes were estimated using the MM/PBSA method. As the entropic contribution was not included, the reported values correspond to the relative binding free energies. Results are summarized in Table 3. Among the four systems, L1 and L3 displayed the most favorable binding free energies, suggesting higher binding affinity to GUCY2C in comparison to L2 and L4. Decomposition of the energetic terms indicated that the more favorable binding observed for L1 and L3 was predominantly driven by stronger electrostatic and van der Waals interactions, whose large negative contributions compensated for their larger polar solvation penalties.

**Table 3:**
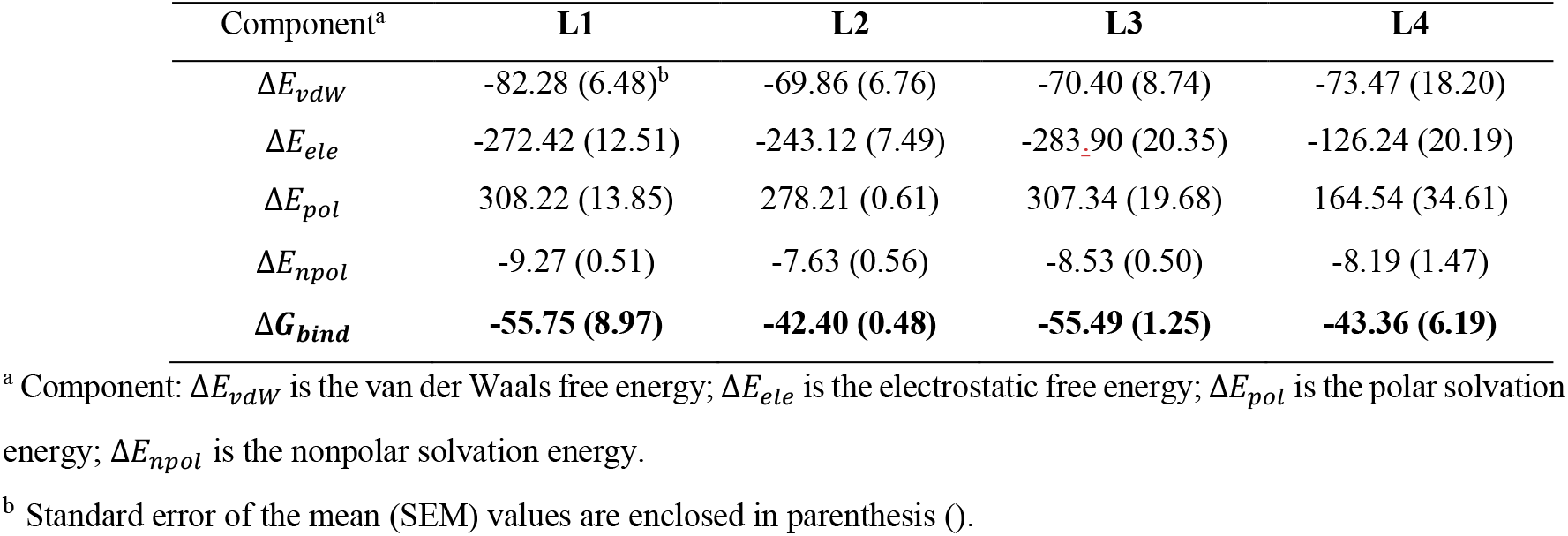
MM/PBSA predicted relative binding free energies of each GUCY2C-scFv complex. All values are expressed in kcal mol-1.

| Component<sup>a</sup> | L1 | L2 | L3 | L4 |
| --- | --- | --- | --- | --- |
| $\Delta E_{vdW}$ | -82.28 (6.48)<sup>b</sup> | -69.86 (6.76) | -70.40 (8.74) | -73.47 (18.20) |
| $\Delta E_{ele}$ | -272.42 (12.51) | -243.12 (7.49) | -283.90 (20.35) | -126.24 (20.19) |
| $\Delta E_{pol}$ | 308.22 (13.85) | 278.21 (0.61) | 307.34 (19.68) | 164.54 (34.61) |
| $\Delta E_{npol}$ | -9.27 (0.51) | -7.63 (0.56) | -8.53 (0.50) | -8.19 (1.47) |
| $\Delta G_{bind}$ | **-55.75 (8.97)** | **-42.40 (0.48)** | **-55.49 (1.25)** | **-43.36 (6.19)** |

<sup>a</sup> Component: $\Delta E_{vdW}$ is the van der Waals free energy; $\Delta E_{ele}$ is the electrostatic free energy; $\Delta E_{pol}$ is the polar solvation energy; $\Delta E_{npol}$ is the nonpolar solvation energy.
<sup>b</sup> Standard error of the mean (SEM) values are enclosed in parenthesis ().

To elucidate the energetic contribution of individual residues of the scFv-GUCY2C interface to the overall binding free energy, per-residue energy decomposition analyses were performed (Figures S5 and S6). L1 and L3 shared highly similar interaction profiles, with dominant contributions arising primarily from V_L_ domain residues Tyr31, Tyr32, and Lys97, and Trp107 from the V_H_ domain. In contrast, L2 exhibited a distinct profile in which contributions from the V_L_ domain were markedly smaller, and binding was instead dominated by a limited number of V_H_ residues. This asymmetry is consistent with the increased structural fluctuations observed for the V_L_ region in its RMSF profile, suggesting that this domain may have been less engaged in the interaction with the receptor compared with the other systems. On the receptor side, Thr76, Leu80, and Leu83 consistently emerged as key contributors across systems. Leu80’s strong contribution is in accordance with previous experimental findings which identified this residue as pivotal in the binding of the anti-GUCY2C arm of PF-07062119 to GUCY2C [16].

### 3.4 Pairwise Interaction Analysis of the scFv-GUCY2C Interface

While both L1 and L3 exhibited the most favorable binding free energies, complex L1 displayed greater structural stability across the simulations and was therefore selected for detailed interfacial analysis. Intermolecular contacts at the L1-scFv-GUCY2C binding interface were monitored throughout the simulations, and the frequency of each pairwise contact was calculated to identify the most persistent interactions.

The interface consists of a mosaic of hydrophobic, polar, and charged interactions. Within the V_L_ domain, Lys97 and Tyr99 formed some of the most persistent interactions with epitope residues. Notably, Lys97 frequently engaged residues in the N-terminal portion of the epitope, while Tyr99 established sustained contacts with Leu80. Although Tyr31 and Tyr32 were identified as key energetic contributors in the MM/PBSA analysis, their contacts were less persistent. Within the V_H_ domain, which includes a larger number of interfacial residues, a correspondingly greater number of residues established high-frequency contacts with the GUCY2C epitope. Among these, Trp107 and Asn59 combined high contact frequency with favorable energetic contributions, indicating a stabilizing role in complex formation.

Importantly, Leu80 emerged as a central interaction hub within the epitope. In addition to its contacts with V_L_ residues, it formed highly persistent hydrophobic interactions with V_H_ residue Trp107, a contact that was maintained throughout nearly the entire simulation and is also observed in the experimental crystal structure. The energetic contribution and frequent contacts formed by Leu80 further support its critical role in antibody recognition.

To further characterize the interface, persistent salt bridges, hydrogen bonds, and hydrophobic contacts were identified using geometric and distance-based criteria (Table S2). Several stabilizing electrostatic interactions were observed, including salt bridges involving Lys97 (V_L_) and Glu104 (V_H_). Additionally, the Leu80-Trp107 hydrophobic interaction was confirmed under more stringent classification criteria. As expected, occupancies defined by these stricter criteria were lower than those obtained from distance-based contact analysis.

## 4. Discussion

Here, a computational workflow was implemented to study the dynamics of the anti-GUCY2C-scFv-GUCY2C interaction and evaluate four alternative scFv designs incorporating different linkers. Across the studied systems, linker choice was found to influence both conformational stability and predicted binding affinity to GUCY2C. L1 and L3 exhibited more favorable binding free energy profiles compared with L2 and L4, while L1 demonstrated enhanced structural stability and greater stabilization of the receptor. Further experimental validation would be required to confirm the computational selection, particularly given that L1 and L3 exhibited comparable relative binding affinities while differing in their effects on receptor conformational stability.

Similar computational work has been previously performed on a ricin-based fusion protein, with different constructs analyzed to study different linkers and active sites [38] and in the design and simulation of a fusion protein comprising the light chain and translocation domain of botulinum toxin A fused to a selected targeting domain as a general framework for fusion protein engineering [39]. These studies assessed the intrinsic stability and structural properties of the engineered constructs in isolation, rather than systematically evaluating their behavior in the context of interaction with their intended molecular target. In contrast, the present study comparatively examined different linker-containing scFv constructs in their bound state to GUCY2C to interrogate how linker composition affects complex stability and antigen binding affinity. An evaluation of the unbound conformational ensembles of each construct would also be valuable, particularly in light of the conformational selection paradigm in antibody antigen recognition, where binding competent states are favored within a pre-existing conformational ensemble [40].

More broadly, it is challenging to correctly determine whether a simulation has reached convergence [41]. ScFvs exhibit conformational transitions spanning multiple timescales, with rearrangements of the CDR loops occurring on microsecond to millisecond timescales, whereas V_L_-V_H_ interdomain dynamics have been reported to occur on much faster timescales, in the low nanosecond range [42]. In this context, it cannot be excluded that slower conformational states relevant to the interaction were not fully sampled in the simulations performed here. Nevertheless, the present analysis focused on the relative behavior of scFv variants within an ensemble of relevant domain orientations potentially involved in antigen binding, which is already accessible using conventional short MD simulations [43]. A more thorough exploration of the conformational landscape of the scFv-GCC system could be achieved in future studies through extended simulations or the application of enhanced sampling techniques.

Collectively, the analyses performed here provided a dynamical insight into the conformational stability and relative strength of interaction between each scFv variant and its target, identifying L1 (GSTSGSGKPGSGEGSTKG) as the most promising linker for the studied anti-GUCY2C scFv. Our results suggest that scFvs containing different linkers exhibit different structural behavior and antigen affinities and illustrate how computational approaches can support rational linker selection and candidate prioritization prior to resource intensive in vitro and in vivo evaluation of molecular function and therapeutic performance.

## ACKNOWLEDGEMENTS

This work was supported by national funds through FCT – Fundação para a Ciência e a Tecnologia under project UID/04349/2025. Computational resources were provided through the FCT Advanced Computing Project 2025.07552.CPCA using the Deucalion Supercomputer.

