## Supplementary material for "Structure-Guided Computational Analysis of Linker effects in an scFv Targeting Guanylyl Cyclase C": Table S1 in Supplementary Information

### Supplementary Tables

**Supplementary Table S1:** MolProbity scores for the 5 best predicted structures of each sccFv model (bold indicates the structure selected for subsequent docking).

| scFv Model | MolProbity Score |
|---|---|
| L1 | S1: 1.20 |
| | **S2: 0.99** |
| | S3: 1.17 |
| | S4: 1.39 |
| | S5: 1.13 |
| L2 | S1: 1.35 |
| | **S2: 1.13** |
| | S3: 1.28 |
| | S4: 1.20 |
| | S5: 1.13 |
| L3 | S1: 1.21 |
| | S2: 1.23 |
| | S3: 1.46 |
| | **S4: 1.19** |
| | S5: 1.41 |
| L4 | **S1: 1.09** |
| | S2: 1.33 |
| | S3: 1.28 |
| | S4: 1.28 |
| | S5: 1.58 |

**Supplementary Table S2:** Equilibrated regions of the MD simulations of each studied system and replica in nanoseconds.

| | R1 | R2 | R3 |
|---|---|---|---|
| **L1** | 100-400 | 50-400 | 100-400 |
| **L2** | 100-400 | 100-400 | 50-400 |
| **L3** | 100-400 | 100-400 | 300-500 |
| **L4** | 100-400 | 100-400 | 50-400 |
| **Unbound GUCY2C** | 75-300 | 100-300 | 100-300 |

### Supplementary Figures:

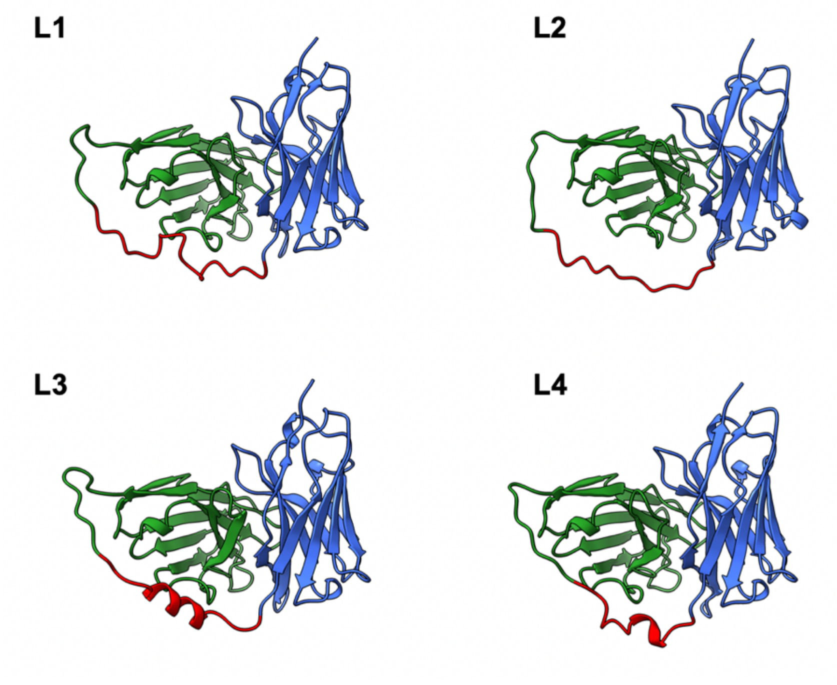


**Supplementary Figure S1:** Best structure for each scFv model (L1-L4) selected based on MolProbity score. The $V_H$ domain is colored in blue, the $V_L$ domain in green and the linker in red.

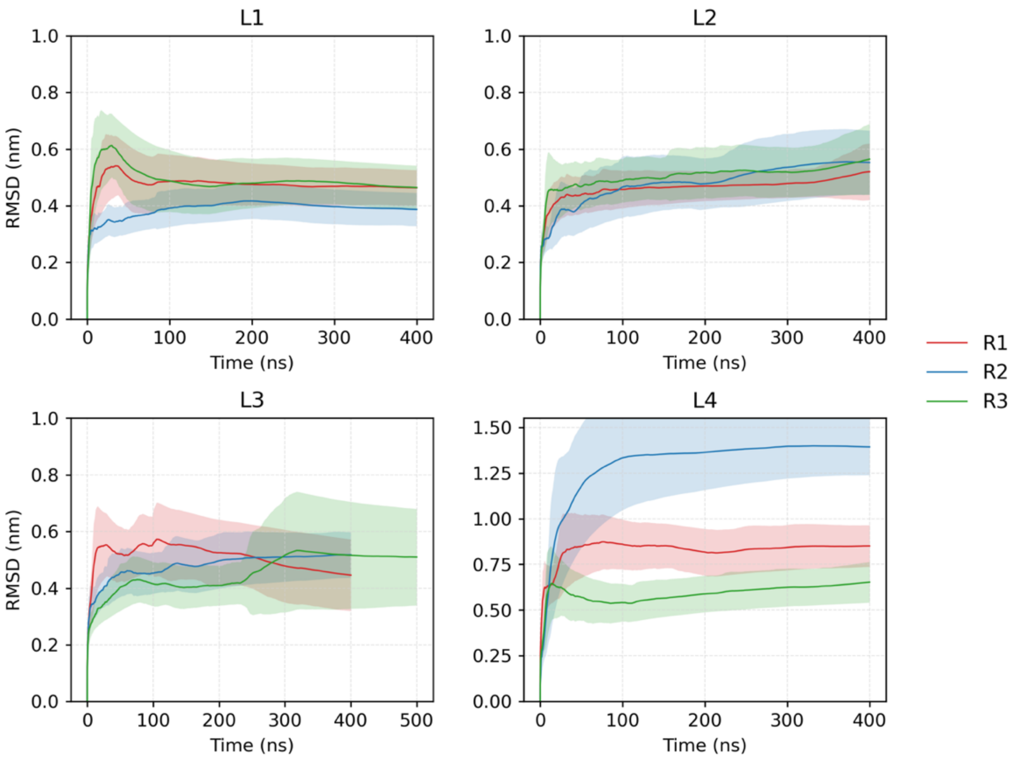


**Supplementary Figure S2:** Average RMSD of the backbone atoms of each scFv-GUCY2C complex (L1-L4), calculated relative to the initial frame of each trajectory, over the course of MD simulations. The three replicates of each system are shown, and the corresponding standard deviations are represented by a shaded envelope.

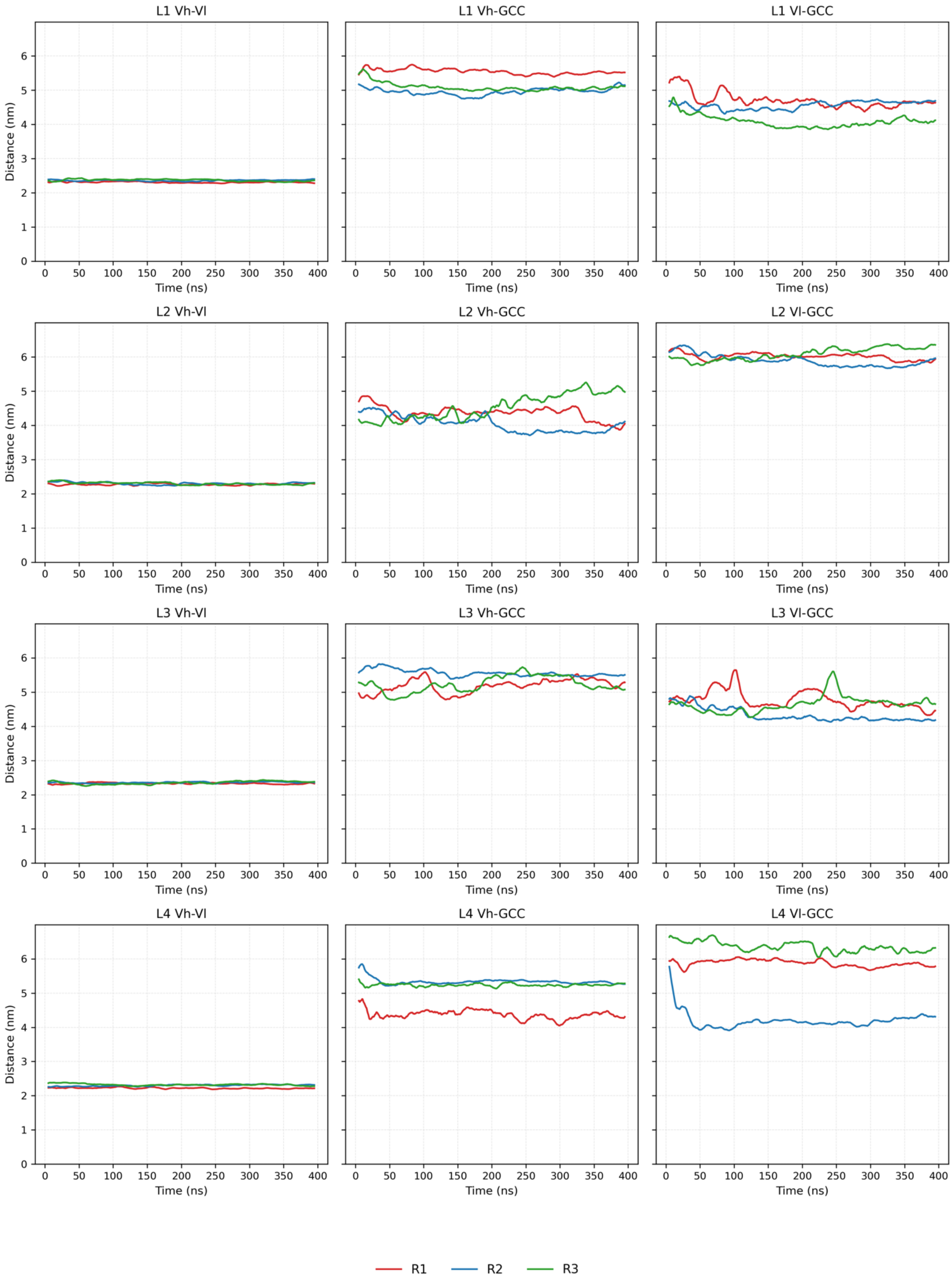


**Supplementary Figure S3:** Time evolution of the interdomain center-of-mass (COM) distances for each scFv-GUCY2C system (L1-L4). Vh refers to the variable heavy domain, Vl to the variable light domain and GUCY2C to guanylyl cyclase C. Data were smoothed using a 10 ns floating average window.

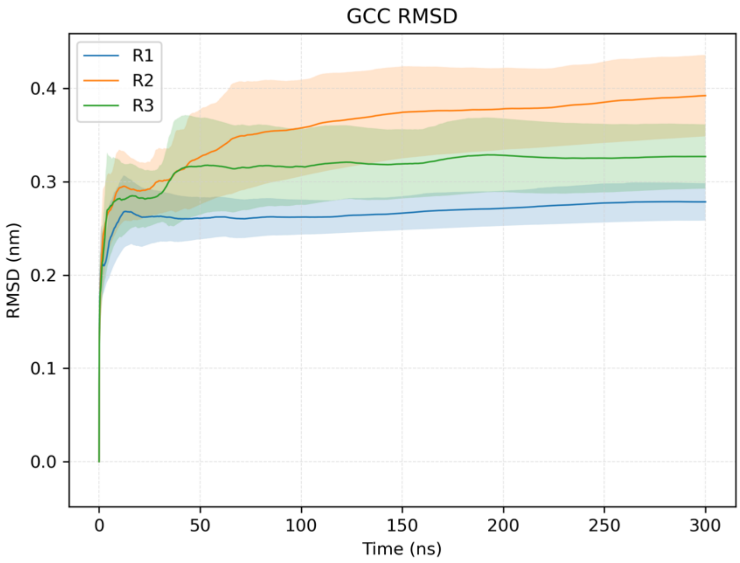


**Supplementary Figure S4:** Average backbone RMSD of the unbound GUCY2C trajectories, calculated relative to the corresponding initial frames. Standard deviations are represented by a shaded envelope.

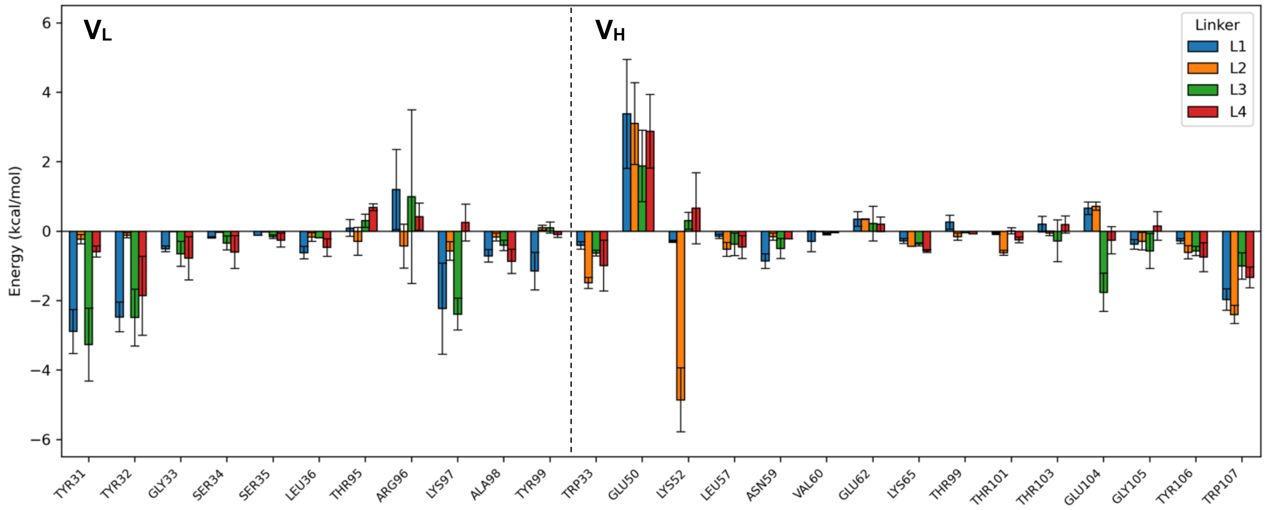


**Supplementary Figure S5:** scFv per-residue contribution to the binding free energy for each scFv-GUCY2C complex (L1-L4). Vertical dashed lines separate residues of the $V_L$ (left) and $V_H$ (right) domains. Values represent the average across replicates, and error bars denote the SEM.

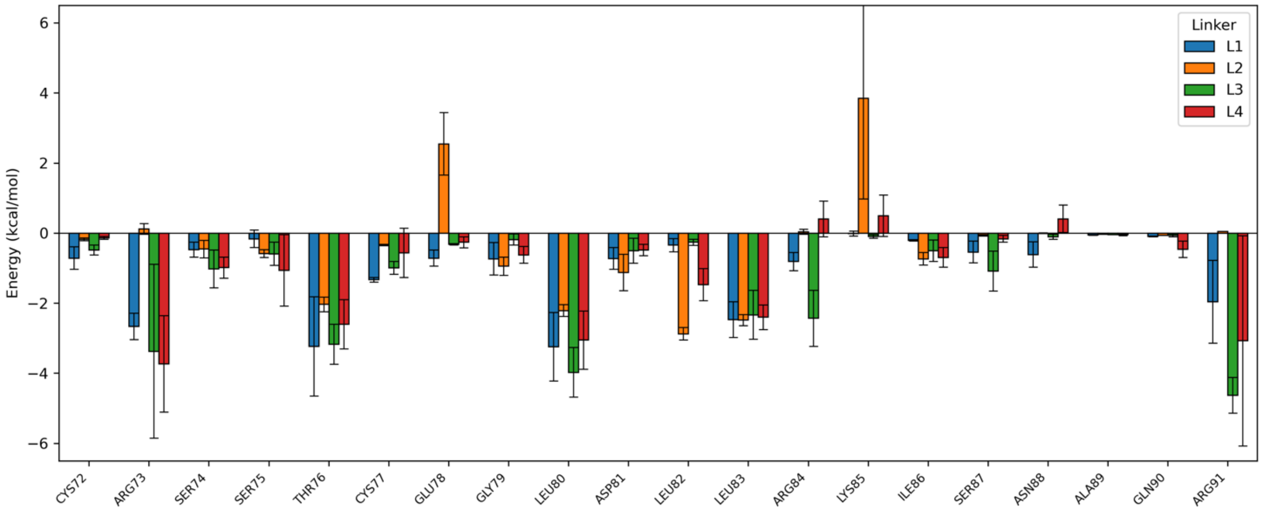


**Supplementary Figure S6:** GUCY2C per-residue contribution to the binding free energy for each scFv-GUCY2C complex (L1-L4). Values represent the average across replicates, and error bars denote the SEM.
